# BioFunctional: A Comprehensive App for Interpreting and Visualizing Functional Analysis of KEGG Pathways and Gene Ontologies

**DOI:** 10.1101/2024.10.08.616405

**Authors:** Alejandro Rodriguez, Antonio Monleon Getino

## Abstract

A comprehensive application designed for the interpretation and visualization of the functional analysis related to KEGG pathways and gene ontologies gives researchers and specialists a tool to get detailed functional information about their data, specifically going deep into biological pathways and gene functions information. By using a variety of techniques and libraries, such as Shiny, htrr, dplyr, tibble, and rvest, we have developed an application that provides a well-designed user-oriented interface with all the facilities to assess their data and start analyzing it directly from scratch through a few steps.

The software allows an exhaustive exploration of KEGG pathways and Gene Ontologies, facilitating the analysis of complex biological processes. To achieve this, functions described in the scripts integrate data manipulation methods and web scraping techniques to extract the necessary information from online official databases, Kyoto Encyclopedia of Genes and Genomes (KEGG) and QuickGo. Furthermore, those functions are computed by parallel processing, resulting in efficient petitions to the database servers and allowing the user to get quick results from a large dataset.

A crucial feature of BioFunctional is its ability to obtain ancestral information for KEGG pathways and gene ontologies, using the techniques described above. This makes it easier to understand the hierarchy of these ontologies and how each sample in a dataset is classified within them, offering users a way to study the dataset at different taxonomic levels directly from the raw data. Additionally, the app implements the capability to create interactive networks, representing all experimental data to visualize the relationships between groups and ontologies without neglecting the established classification. This is a primary tool for understanding the meaning of the relationships observed within the displayed system.

**Key features of Biofunctional include:**

- Interactive visualization: Create and explore networks to visualize relationships between groups and ontologies.
- Hierarchical analysis: Trace ancestral information for KEGG pathways and Gene Ontologies to understand hierarchical classifications.
- Efficient processing: Employ parallel processing for rapid data analysis, even with large datasets.
- Intuitive interface: A user-friendly design simplifies data exploration and analysis.

Due to these attributes, the software represents a valuable tool for analysts involved in the study of KEGG pathways and Gene Ontologies. By providing an intuitive interface with advanced data processing techniques, it empowers researchers to unravel the intricacies of biological functions and gain insights into the relationships between genes or molecular components.

**Supplementary information:** https://github.com/alexrodriguezmena/BIOFunctional

## 1 Introduction

In biology, functional analysis is a start-point procedure used to understand the functions and interactions of genes and proteins that happen in organisms [11]. It includes a sort of techniques to interpret proteomic and genomic data, providing insights into the basic mechanisms regulating physiological and cellular processes [14].

The “foundation” of functional analysis falls to the Gene Ontology (GO) project, which offers a dictionary for defining genes and proteins according to their molecular functions, biological processes, and cellular components [12].

Comparative genomics and evolutionary studies have become simpler by the systematic classification of gene and protein properties, which provides researchers with insights into the regulatory processes and functional roles of these across species [13].

Another essential component of functional analysis is pathway analysis, which focuses on finding and characterizing biological pathways that are enriched with proteins or genes that express themselves uniquely [21]. These pathways are complex networks of molecular interactions that regulate a variety of biological functions, so researchers can use them to clarify the functional context of genes and proteins within these pathways.

By identifying biological terms overrepresented in a set of genes or proteins of interest, functional enrichment analysis works as a complementary tool to pathway analysis [18]. This method consists in statistically evaluating gene or protein annotations against a background reference, guiding the formulation of predictions and the validation of experimental results by returning information about the biological importance of gene or protein sets, and letting researchers find enriched functional terms associated with specific biological processes, molecular functions, or cellular components.

Gene Ontologies (GO) are a set of terms and relationships that describe the biological characteristics of genes and proteins in standardized and structured terms [12]. Those ontologies are classified in three different domains, as was commented, molecular functions, biological processes, and cellular locations associated with genes and proteins in different organisms.

A molecular function (MF) ontology describes the molecular-level activities or functions performed by a gene product without giving evidence of where or when this activity occurs. MF terms include, inter alia, “catalytic activity” and “DNA binding” [10].

The biological process (BP) ontologies represent the accomplishment of a biological program, which consists of a set of steps to achieve a main goal; this can be carried out by a specific collection of molecular functions performed by specific genes, as explained above [8]. BP terms include, inter alia, “apoptosis” and “cellular cycle” [17].

Cellular component (CC) ontologies are all the cellular locations where molecules are findable and the molecular functions are performed by those [17]. CC terms are ‘organized’ in three types of components: the cellular anatomical entity, in which we find the plasma membrane; the virion component, where the viral proteins take action; and the protein-containing complex, where gene products, such as protein-lipid complex, are located.

Therefore, those three main domains have nothing in common as each of them represent different things, but can those be somehow connected, that’s what we are going to discover [16].

Structure of each GO term can be described in a graphical way, where each node represents a gene ontology, and the relationship between two or more nodes are edges (Supplementary material Figure 1 & 2) [17]. GO doesn’t follow a clear hierarchy, the closest thing to a contrasted classification are the three main domains that encompass all the ontologies. So it basically shows a kind of parent/child relationship between all the nodes, where the children are more specific functions each level and the parents represent the main subgroups term that represent a bunch of ontologies [16]. Although this parent/child relationship is not as simple as it can seem, as there are different types of relations that can happen betwixt the ontologies, being the main ones: *is a*, means that one node is a subtype of another; *part of*, means that one ontology is necessarily part of another one, but only one way, so for example if we are talking about mitochondrial (cellular component) we will always relate it to cytoplasm and it’s known that without it the mitochondria can exists, but if the study is centered in the cytoplasm it is not essential or necessary to talk about the mitochondria; *has part*, has the same logic as *‘part of’* but instead of a node being a part of another, it means a node has another one as a part of it; and *regulates*, means when a process or a function directly affects the performance of a distinct one [21]. So why are we not able to have a classification if it is just a simple graph with a set of different relations between nodes at distinct levels; because not all the relationships allow grouping, as for example regulating one ontology doesn’t mean to be part of it or to take part in the performing of this activity, hence that’s the problematic point, as it’s really complex to create a taxonomy when not all the relations are truly significant in order to store it in a category [20].

So replying to the main question announced before, all the ontologies can have more than one parent by diverse connections, noting all along the graph that the different relationship permit an ontology to be a subtype of molecular functions, while regulating biological processes, which means that all the ontologies can be somehow connected even the three big domains [21].

In addition, GO are in constant movement, as are constantly revised and updated in order to achieve the main objective; describe and represent all the biological knowledge [12].

KEGG (Kyoto Encyclopedia of Genes and Genomes) is a sort of databases that stores information about biological functions and the metabolic processes of all the organisms, standing out above the others, the KEGG pathways database shows the molecular interactions, reactions and relations that happen in a organism [13].

KEGG pathways are a collection of pathway maps that represent molecular interaction between genes or proteins [20]. These diagrams represent the relationships between molecules and how they are connected or regulated, helping to understand the biological function of each of them.

Something to highlight is the hierarchy of the KEGG pathways database, which stores all the information classified in three domains: the most general group organizes the pathways in categories by the function it has, so some examples could be Metabolism or Cellular processes, then second domain tries to specify a bit more in the classification of the pathways, dividing them in organism for metabolism or in the different processes a cell can perform, and for the last group all the metabolic pathways are described as explained above [13].

Also, this database is strongly connected with several distinct databases, giving the opportunity to the researcher to get all the information needed for the study of one biological function or pathways in the same platform [13]. Hence, some of the databases referenced are NCBI, Uniprot or GenBank, giving the chance to access information about the genes or proteins of interest.

## 2 Objectives

The aim of this project is to develop an user-friendly application to interpret and visualize the functional analysis for KEGG pathways and Gene Ontologies from an experimental dataset, looking for exhaustive insights about the biological pathways and genes, so the researchers have a tool to investigate and get deeper in molecular biology and bioinformatics.

## 3 Methods

### 3.1 Dataset

The main reason for the development of this application was the necessity to do a comprehensive functional analysis for the dataset resultant of an experimental analysis. *(Supplementary material. Figure 1 & 2)*

### 3.2 Data Pre-Processing

Raw datasets were full of useless observations, combining groups of different samples or instances without values, so before starting processing the data some filters were applied in order to delete those observations that would cause errors or false results when the application gave an output.

Filters are added so the dataset can be adapted to the algorithm and get rid of those columns that won’t be used. Also, a new category was added in the filter feature of the software, and it was the possibility of adding an extra column in the table named “Disease” (defined by our dataset sample case), where the diseases name would be extracted from the raw data and stored for further analysis, only useful when your file stores experimental data of more than one disease, basically it request the input column to inspect and the names of the different diseases.

#### 3.2.1 Differential expression analysis

Differential expression analysis is a method used in fields like genomics, transcriptomics, and proteomics, which compares two or more experimental groups, like healthy states with disease in different stages, by identifying statistical significant differences between the biological measures of each observation, being expression levels, concentrations… some examples of possible measures used.

The aim is to find which observations between groups are significantly regulated, requiring a few statistical techniques or processes [2][22]. In summary, it involves modeling read counts using a negative binomial distribution [Figure 1] [23].

**Figure 1.**
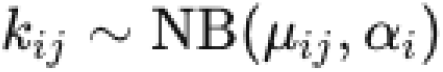
Negative binomial distribution. where: μij is the mean count for gene i in sample j, αi\alpha_iαi is the dispersion parameter for gene iii.

Followed by the estimation of the effects of experimental groups conditions using a log-linear model [Figure 2].

**Figure 2.**
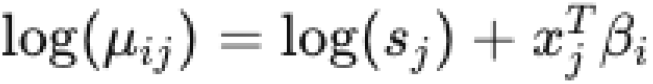
Log linear model equation. where: log(s_j) is the log of the size factor for sample j, x_j is a vector of covariates (e.g., condition labels) for sample j, β_i is a vector of coefficients for gene i.

Proceeding testing for differential expression with hypothesis testing, using a likelihood ratio test [Figure 3]

**Figure 3.**
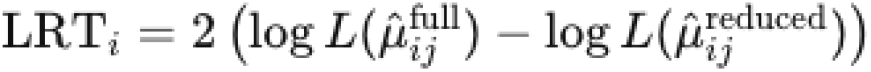
Likelihood ratio test. where: L(μ^ijfull) is the likelihood of the data under the full model; L(μ^ijreduced) is the likelihood under the reduced model

Ending with a multiple testing correction to comprove the False Discovery Rate (FDR) [24] of each observation.

#### 3.2.2 GSEA

Gene Set Enrichment Analysis (GSEA) [Figure 5] is a computational method used to determine a set of genes, or gene ontologies in this case, that happen to be statistically significant in a two group comparison between biological states in a sample [18]. Despite the differential analysis, in this case collective behavior of genes sharing common biological functions pathways are compared by following a few steps to compute the enrichment score [Figure 4]. First step is ranking genes based on a metric computed before-hand, it can be the p-value computed during the differential expression analysis; then gen sets have to be established based on a user-choice theme, it can be biological functions or pathways; and finishing the process with enrichment score mentioned before, which measures how much overrepresented is a gene of a ranked gene set list [25].

**Figure 4.**
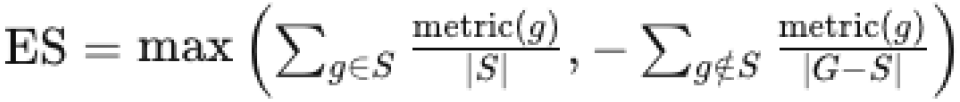
Enrichment analysis value. where: metric(g) is the score of gene g; |S| is the number of genes in set S; |G-S| is the number of genes not in S.

**Figure 5.**
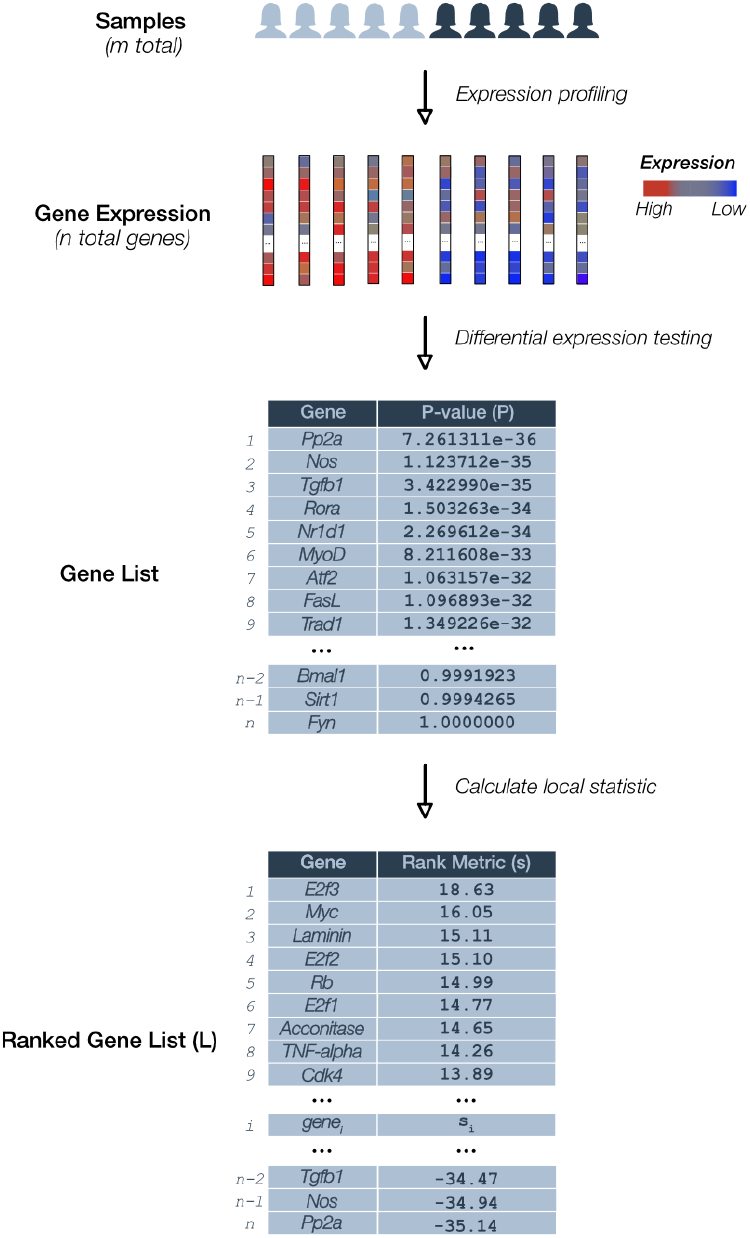
Gene Set Enrichment Analysis general scheme. This figure illustrates the results of a gene enrichment analysis, which identifies sets of overrepresented genes in a dataset to reveal significant biological processes or pathways. The top section shows two groups of samples under different conditions. Heatmaps display gene expression levels, with red indicating high expression and blue indicating low expression. Tables list genes by P-value and rank metric, highlighting their statistical significance and contribution to the enrichment score. This analysis helps identify key genes and pathways that differentiate the experimental conditions, enhancing our understanding of the underlying biological mechanisms.

**Figure 6.**
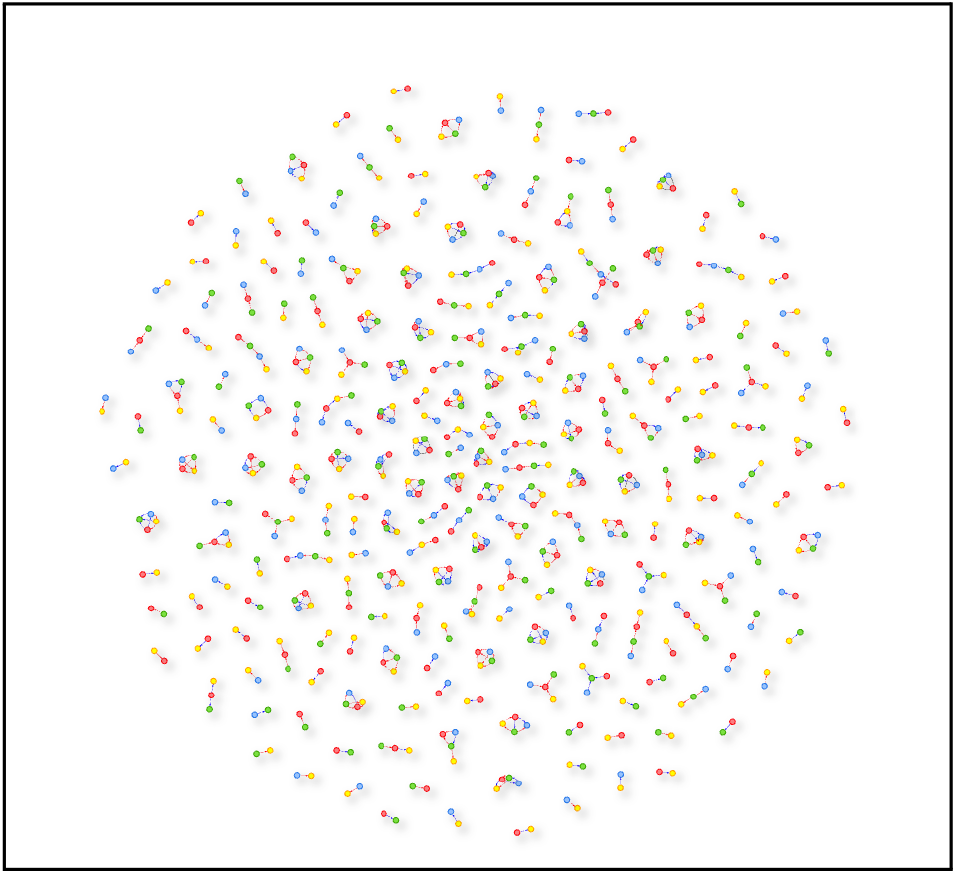
GO network representation. In the network representation for gene ontologies, nodes related are configured to represent the biological entities, as the genes or the ontologies would be, and the edges represent the relationships between them, in this case edges connect the same ontology represented in a different experimental group (coloured to differentiate). Also the edges represent how the relations can be, black line represents that the relation between two groups is neutral, the red line represents that the first group is overexpressed in comparison to the other and the blue line represents the reverse situation.

**Figure 7:**
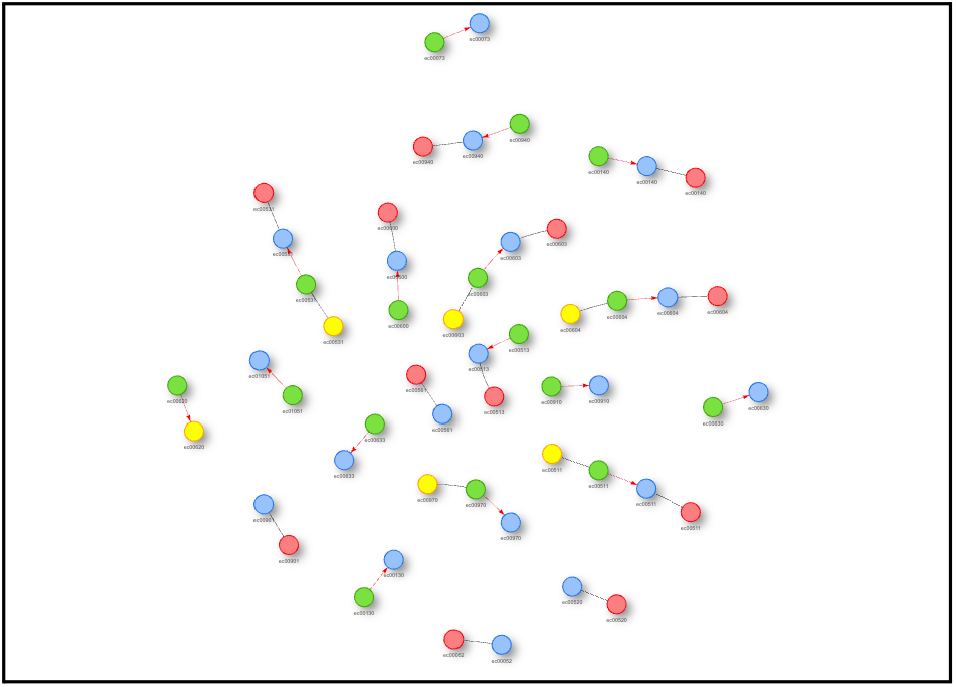
KEGG Pathway network representation. In the network representation for KEGG pathways, nodes related are configured to represent the biological entities, as the genes or the ontologies would be, and the edges represent the relationships between them, in this case edges connect the same ontology represented in a different experimental group (coloured to differentiate). Also the edges represent how the relations can be, black line represents that the relation between two groups is neutral, the red line represents that the first group is overexpressed in comparison to the other and the blue line represents the reverse situation.

#### 3.2.3 KEGG ancestry algorithm

Once the KEGG data is filtered and ordered, it’s ready for web scraping, hence this technique is used for extracting information from a website [9][20].

In this case, the function will extract data from the online database Kyoto Encyclopedia of Genes and Genomes and store it in our initial file by using the instances found in the column that defines the KEGG pathway IDs.

In particular, *rvest* library in R was used to do the web scraping, it allows the user to inspect, analyze and extract the html content from the chosen webpage [13] [26]. Therefore, by the use of *read_html()*, the database was inspected and once there, the function *html_table()* was used to extract the info needed to get the taxonomy for each iteration.

Once data is extracted, the function defined transforms and adapts the data to an optimal format so it can be added to the main and original dataset, as the main domain column and the subdomain column instances for each pathway.

All the taxonomy of each ontology is noted in the dataset and ready to be represented and analyzed in further actions([*Supplementary material. Code 1)*.

#### 3.2.4 GO ancestry algorithms

After the data is filtered and ordered as needed, the taxonomy or classification of gene ontologies process can go ahead [3][8][11][17]. As was explained before, GO hasn’t a defined hierarchy and grouping all of them in different levels requires a too complex procedure, so let’s start by doing secure steps and creating a classification methodology from the most general terms [27]. Overall, what will be done is relating all the ontologies in the file to the very next level from the three big domains so we can get more specific information about these [4].

Firstly, all the ancestors of each of the ontologies given have to be collected and added to the dataset [12] [19], so a function to get the ancestors is defined to send a HTTP request to the Quick Go API and collect the information specified, then the data obtained is comproved to delete those ontologies that are obsolete and not studiable anymore [28]. Once it is ready, is added and returned in an “ancestors” column on the original dataset.

Then, in order to get one level down from the main domains is needed to get all the children from them so a new level of specificity is established and our ontologies dataset can be classified in a more accurate way. Therefore, following the same logic as the algorithm to get the ancestors of all data ontologies, a request to the Quick Go API is done, and the children information of the ontology selected (MF, BP or CC) are stored in json format ready for the next step.

Before the main algorithm, it is needed to create a function to treat the data and extract only the information that will be used in the classification creation as the name and ID of each ontology.

Last step is to connect our data ontologies with the second taxonomy level defined, so a function has been declared in which all the ancestors of each domain are thrown to the dataset of children of their main domain, checked by looking at the *ont_description* column, looking for a coincidence between the ancestors and the children, that will mean a connection, once the bond is done the child correlated past to be the main ancestor of our ontology of interest, and the ID and name of each is stored in the original data file (*Supplementary material. Code 2 & 3)*.

### 3.3 Network Visualization

The aim of the network visualization was to create an interactive network where all the experimental and biological relationships could be displayed, and let the user interpret the results in a simple manner [5][21].

So, library visNetwork from R is essential, as it allows to configure the network as the data demands and the user requires [29].

Filters are added to the network configuration, and can be edited by the user as needed, looking to represent just the data of interest and separate observations that would be senseless to display in the same interactive graph (*Supplementary material. Code 4)*

Hence, the aim of this network representation is to see, by setting some conditions, whether an ontology is represented or not.

### 3.4 App Interface

The idea was to create an intuitive software so any user can use it, without having any insight of biology, bioinformatics, statistics or any other subject related to it, so to do that *shiny* was the library to use [1].

*Shiny* is a R library that allows the user to run an interactive web app directly from the R environment without any knowledge of HTML, CSS or any other web programming language [6], which is good point for the programmer as a lot of time is saved and for the user as it creates a extremely instinctive server [30]. But by the time, this package has evolved and transformed into a source of packages that permits you to add a lot of features to your code.

One of those packages is shiny.dashboard [31], which gives an extra layer of organization and design to a basic shiny app, facilitating the interpretation and analysis of the data by a user, presenting it in a concise and clear manner [7].

### 3.5 Parallelization

Parallelization of the code involves the execution of multiple tasks simultaneously, which reduces the computation time and improves the performance of the functions; when functions have to iterate on each observation of a large dataset, as in this dataset case, implementing this technique is crucial to get an efficient compilation of the results.

In this case, a tool for parallelization is *future*.*apply* library in R, which allows a simple execution of functions in parallel. The main benefit of using it, as obvious, is the reduction of execution time by using a precoded library so everybody can do parallelization, even not having any knowledge on it, but it also presents a really good scalability as it is adaptable to any computing system, which relapse in the use of it for every kind of user. Although the use of a library like future.apply, which does the parallelization process for the user, presents some limitations: the user has no control on how this process is implemented and can’t optimize as his code requires, blocking the possibility to adjust the parallel processes as wanted by the user.

For the software, it was easily implemented by the *future*.*lapply()* function, after setting the configuration of the process by the use of the *plan()* function [32].

### 3.5 AI prompt interpreter

As known several companies have been creating conversational chat AIs since artificial intelligence appeared in our lives, some examples are OpenAI, copilot, gemini… [33]

All of them suppose an effective way of interpreting different inputs, and we will take advantage of them as we need an interpretation of the results for non-scientific users.

Basically, several prompt interpretation messages for gene ontologies functional analysis will be written until a suitable answer is returned, so all our users can get the requested conclusions for their project.

## 4 Results and Discussion

### 4.1 Data preprocessing

The software is completely able to retrieve information from online databases, like Kyoto Encyclopedia of Genes and Genomes (KEGG) and QuickGo by using the methods explained previously, being also able to construct usable and optimal dataframes for further processes from the data obtained.

Also a classic functional analysis was done so we can compare the effectiveness and results of our software with a classic approach of a dataset *(Github. analsis_functional)*

### 4.2 Hierarchical Analysis

A preliminar hierarchical analysis structure has been determined for both, KEGG and GO, giving the chance to the users to delve into all the taxonomy levels determined. On the one hand, KEGG pathway have been easier to classify, as there was a clear structure determined beforehand, but on the other hand, gene ontologies hierarchy is still in development because, as it was explained, taxonomy is not defined beyond the top three ontologies, from where a parents-child relation is created between

all the other ontologies, which makes challenging to assemble GO in different groups of complexity; but first steps have been done, and first level of taxonomy from the main three ontologies has been established and proven, achieving enthusiastic results, showing that gene ontologies can be classified in more classes.

### 4.3 Interactive Network

Interactive network has been developed in order to interpret and visualize KEGG pathways and Gene Ontologies relationships (*Supplementary Figure 3 & 4)*.

After several trials on distinct configurations, an interpretable network for both cases have been displayed, adjusting the filters and code for both of them as explained before.

The “select by” feature has been key on the classification of the ontologies and allows the user to select the nodes of interest and reorganize them.

### 4.4 Scalability & Performance

Initial tests affirmed the effectiveness of the algorithm when working with large datasets, concluding that the application is scalable and can run any dataset size without any problem. The parallel processing techniques implemented allows the web scraping algorithms to work quicker, optimizing their performance and getting results five times faster.

### 4.5 AI prompt Interpreter

Once several trials were done, a sample prompt for AIs like chatgpt could interpret the results and give a clear answer to know how the most expressed gene ontologies between groups were behaving and how to behave after this analysis *(Supplementary material. Text 1*).

## 5 Conclusions

Objectives have been accomplished, as a user-friendly app has been developed for the interpretation and representation of functional analysis of gene ontologies, as researchers can use it to understand hierarchically how their data is related and so on which biological functions and molecular relations are crucial in the dataset.

Otherwise, by comparing the classical functional analysis computed with the software developed, a huge difference is noted as the classic approach suppose a lot of time invested in understanding your dataset and how it can be related with lots of algorithms to be programmed so you can get suitable plots that can explain how your data behave. Meanwhile, a user can get all the answers for all the questions he can have by using BIOFunctional in a few minutes, with more complex and understandable networks for all the different samples in real time filtering. Then, it’s easier to justify the effectiveness and good interpretation experience the app gives to the users.

Moreover, the AI study shows that official conversational artificial intelligence created by different developers can output useful conclusions of what is happening in the case studied, all by submitting the prompt created input that gives AI the information needed to give a non-expert user the keys to deeply understand what happens in his data.

In conclusion, high-throughput biological data can be understood by any kind of researcher or company by getting the most relevant biological pathways in the dataset information processed along the study of a data case.

## Acknowledgement

First and foremost, I would like to thank Antonio Monleon for supervising this project, as he always was there to help me to decide which were the right decisions in all the steps of the project, and also for all the knowledge he transferred to me, which for sure will be useful in all my career. Then I want to mention Albert Martin, who was also at UB Biost3 group and participated in the first steps of the project, giving me some insights of how to approach the network representation and solved lots of problems we had. Finally, to the entire team of the Sant Joan de Deu Hospital of the PI17/01540 project and to Javier Méndez; as the data used in this analysis were derived from a publicly accessible dataset generated as part of the project “PI17/01540”, funded by the Instituto de Salud Carlos III. All individual-level data have been anonymized to protect participant privacy [34].

## Abbreviations

GO: Gene Ontology
MF: Molecular Function
BP: Biological Process
CC: Cellular component
KEGG: Kyoto Encyclopedia of Genes and Genomes
GSEA: Gene Set Enrichment Analysis.

