## Supplementary material for "BioFunctional: A Comprehensive App for Interpreting and Visualizing Functional Analysis of KEGG Pathways and Gene Ontologies"

The data used in this analysis were derived from a publicly accessible dataset generated as part of the project "PI17/01540", funded by the Instituto de Salud Carlos III. All individual-level data have been anonymized to protect participant privacy.

**OUTLINE:**


**Supplementary Table 1:** KEGG Pathway dataset.
**Supplementary Table 2:** GO dataset.
**Supplementary Figure 1:** GO hierarchy.
**Supplementary Figure 2:** GO hierarchy 2.
**Supplementary Text 1:** IA prompt text to get conclusions about the dataset case.
**Supplementary Code 1:** R script function to get hierarchical order of KEGG Pathways by requesting to genome.jp database
**Supplementary Code 2:** R script function to get ancestors of a Gene Ontology by requesting to Quick Go API.
**Supplementary Code 3:** R script function to get on extra level of hierarchy in gene ontologies classification.
**Supplementary Code 4:** R script functions to configure the network edges color and direction.

| ONTOLOGY | ONT_DESCRIPTION | FDR | EA_VALUE | sample | GROUP | GROUP_1 | GROUP_2 | UP_DOWN | Disease |
|---|---|---|---|---|---|---|---|---|---|
| ec00901 | Indole alkaloid biosynthesis | 0 | 72.9743935309973 | FECAL | F.EC.3M vs F.EC.6M | EC.3.meses | EC.6.meses | NEUTRAL | EC |
| ec00901 | Indole alkaloid biosynthesis | 0 | 72.9743935309973 | FECAL | F.EC.3M vs F.EC.6M | EC.3.meses | EC.6.meses | NEUTRAL | EC |
| ec00901 | Indole alkaloid biosynthesis | 0 | 60.8849325337331 | FECAL | F.C vs F.CU.3M | control | CU.3.meses | NEUTRAL | CU |
| ec00901 | Indole alkaloid biosynthesis | 0 | 60.8849325337331 | FECAL | F.C vs F.CU.3M | control | CU.3.meses | NEUTRAL | CU |
| ec00073 | Cutin, suberine and wax biosynthesis | 0.08408182821641 86 | 6.61620234604106 | FECAL | F.C vs F.CU.DE | control | CU.debut | DOWN | CU |
| ec00140 | Steroid hormone biosynthesis | 2.86025171801287 e-13 | 6.13392977018953 | FECAL | F.C vs F.CU.3M | control | CU.3.meses | NEUTRAL | CU |
| ec00140 | Steroid hormone biosynthesis | 2.86025171801287 e-13 | 6.13392977018953 | FECAL | F.C vs F.CU.3M | control | CU.3.meses | NEUTRAL | CU |
| ec00901 | Indole alkaloid biosynthesis | 0 | 53.3467980295567 | FECAL | NA | control | EC.6.meses | NEUTRAL | EC |
| ec00901 | Indole alkaloid biosynthesis | 0 | 53.3467980295567 | FECAL | NA | control | EC.6.meses | NEUTRAL | EC |

**Supplementary Table 1: KEGG Pathway dataset.** 10 highest enrichment value KEGG Pathway dataset of the total 152 observations, where *ONTOLOGY* column shows the ontologies IDs, *ONT_Description* stores the ontologies names for KEGG, *SAMPLE* refers on how the observation was obtained, *GROUP* refers to the experimental groups comparison (*GROUP_1* and *GROUP_2*) of each instance (ex. control.vs.3.months), *EA_VALUE* is the enrichment analysis resultant value, and *KEGG_up_DOWN* describes which of the groups in the comparison regulates the other by the enrichment value specified, for instance having the group comparison of *control vs 3 month*, if our variable value is *down-regulated* it means it is regulated by the control observation and the other way around if it is *up-regulated*.

| ONTOLOGY | ONT_DESCRIPTION | FDR | EA_VALUE | sample | GROUP | GROUP_1 | GROUP_2 | UP_DOWN | Disease |
|---|---|---|---|---|---|---|---|---|---|
| GO:0009381 | molecular_function | 3.78417207147486e-06 | 9.99143496936184e-05 | FECAL | fecal.EC.3.meses_vs_fecal.EC.6.meses | EC.3.meses | EC.6.meses | DOWN | EC |
| GO:0043190 | cellular_component | 3.58764855923344e-06 | 9.93515592386268e-05 | ORAL | oral.CU.6.meses_vs_oral.CU.debut | CU.6.meses | CU.debut | DOWN | CU |
| GO:0042953 | biological_process | 3.62276403853556e-06 | 9.93515592386268e-05 | ORAL | oral.CU.6.meses_vs_oral.CU.debut | CU.6.meses | CU.debut | DOWN | CU |
| GO:0016282 | cellular_component | 5.17642924269257e-06 | 9.93476227732151e-05 | ORAL | oral.Control.control_vs_oral.CU.6.meses | control | CU.6.meses | UP | CU |
| GO:0033290 | cellular_component | 5.17642924269257e-06 | 9.93476227732151e-05 | ORAL | oral.Control.control_vs_oral.CU.6.meses | control | CU.6.meses | UP | CU |
| GO:0004650 | molecular_function | 1.38777878078145e-14 | 9.87736462669229e-13 | FECAL | fecal.EC.3.meses_vs_fecal.EC.6.meses | EC.3.meses | EC.6.meses | DOWN | EC |
| GO:0051539 | molecular_function | 2.08177258764763e-07 | 9.82776851602788e-06 | FECAL | fecal.Control.control_vs_fecal.CU.3.meses | control | CU.3.meses | UP | CU |
| GO:0043139 | molecular_function | 2.10595039629169e-07 | 9.82776851602788e-06 | FECAL | fecal.Control.control_vs_fecal.CU.3.meses | control | CU.3.meses | UP | CU |
| GO:0009097 | biological_process | 3.3137827593066e-08 | 9.81826491828843e-07 | FECAL | fecal.CU.3.meses_vs_fecal.CU.debut | CU.3.meses | CU.debut | DOWN | CU |

**Supplementary Table 2: GO dataset.** 10 highest enrichment value GO dataset of the 5000 total observations, where *ONTOLOGY* column shows the ontologies IDs, *ONT_Description stores the main domain name for GO (molecular function, biological process or cellular component), SAMPLE* refers on how the observation was obtained, *GROUP* refers to the experimental groups comparison (*GROUP_1* and *GROUP_2*) of each instance (ex. control.vs.3.months), *EA_VALUE* is the enrichment analysis resultant value, and *GOEA_up_DOWN* describes which of the groups in the comparison regulates the other by the enrichment value specified, for instance having the group comparison of *control vs 3 month*, if our variable value is *down-regulated* it means it is regulated by the control observation and the other way around if it is *up-regulated*.

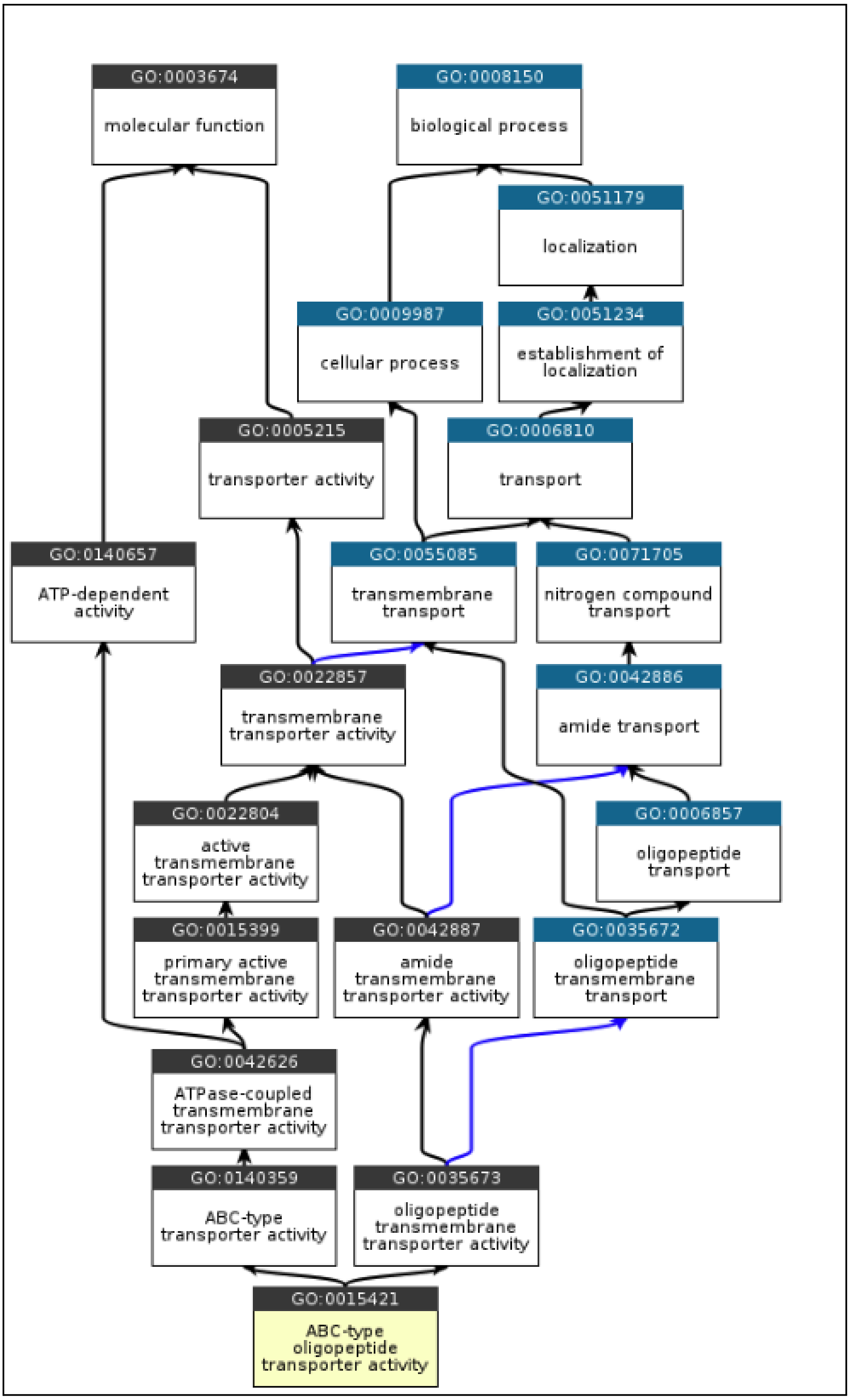


**Supplementary Figure 1: GO hierarchy.** Sample of GO hierarchy tree, where can be checked how arbitrarily the structure of it is and the relationship between the "nodes".

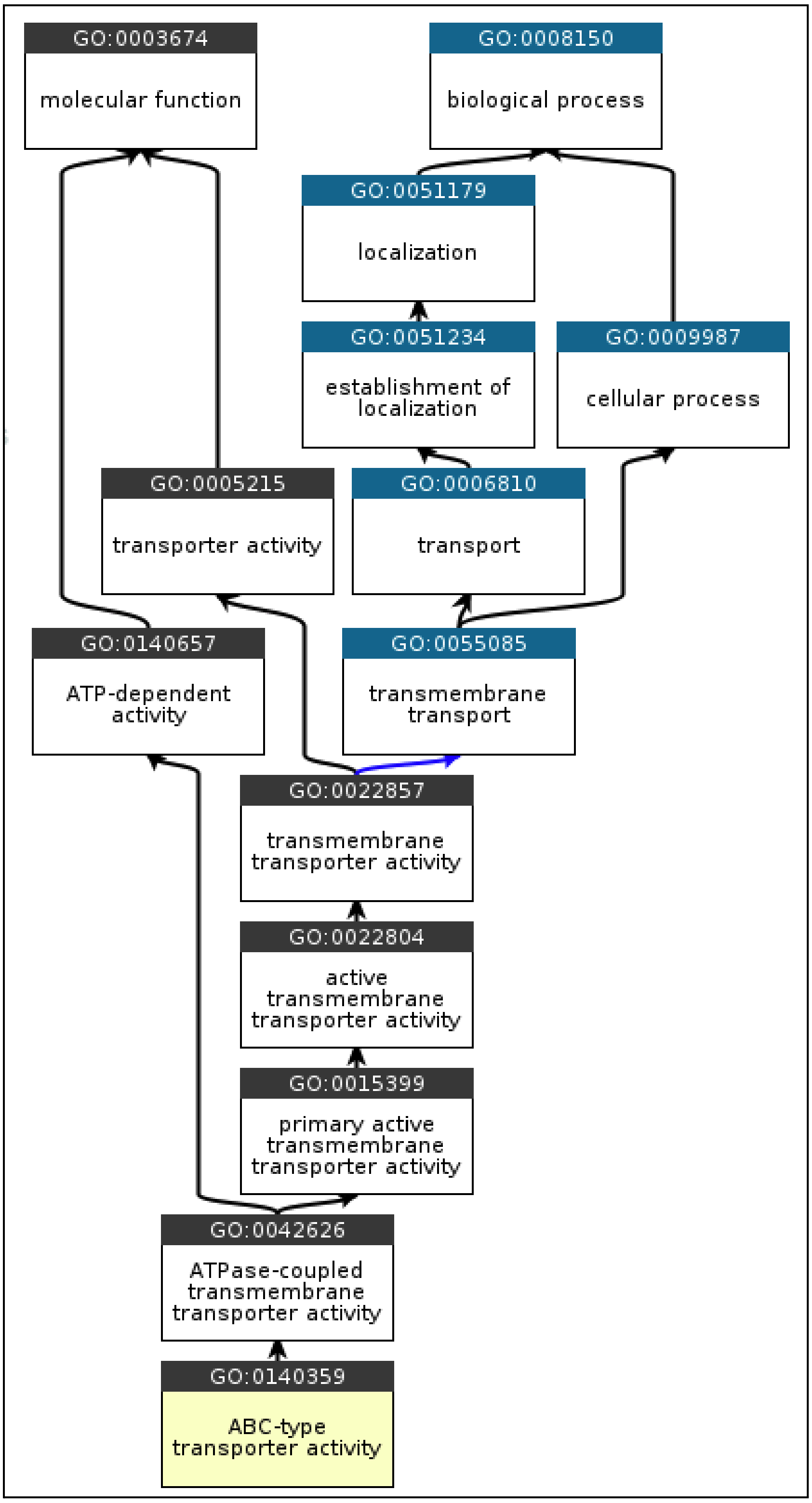


**Supplementary Figure 2: GO hierarchy.** Sample of GO hierarchy tree, where can be checked how arbitrarily the structure of it is and the relationship between the “nodes”.

*‘Write an exhaustive analysis focusing on biological experimental conclusions to learn how the diseases are doing on a Gene Ontology (GO) enrichment dataset for CU or EC, taking the main ideas for all the dataset without specifying in each of them. The dataset includes enrichment information for the 15 most enriched ontologies in the dataset. Each entry in the dataset has the following attributes*

*Ontology: The GO identifier.*

*Sample: Sample used for enrichment analysis .*

*Description: Description of the GO term.*

*Disease: Disease studied.*

*Group_1: First group for comparison.*

*Group_2: Second group for comparison.*

*ea_value: Enrichment value.*

*first_ancestor: The most general ancestor node in the GO hierarchy.*

*So here are the gene ontologies to analyze: ’*

**Supplementary Text 1: IA prompt text to get conclusions about the dataset case.**

```r
# Define la función para procesar los datos de KEGG y devolverlos
ancestors_kegg <- function(data) {
  relations <- unique(data$ONTOLOGY)
  processed_data <- data.frame()  # Inicializa un dataframe vacío

  for (i in seq_along(relations)) {
    relation <- relations[i]

    # Obtener las reacciones e interacciones de la página web
    metabolic_domain <- ""
    metabolic_subdomain <- ""

    # Leer la página web y extraer los datos de la tabla
    url <- paste0("https://www.genome.jp/dbget-bin/www_bget?pathway:", relation)
    webpage <- read_html(url)

    # Realizar el web scraping para obtener la tabla
    tables <- html_nodes(webpage, "table")
    table <- html_table(tables[[1]], fill = TRUE)

    # Encontrar la fila que contiene la información relevante
    class_row_index <- which(table[, 1] == "Class")
    if (length(class_row_index) > 0) {
      class_row <- table[class_row_index, ]

      # Obtener el valor de interacción y reacción de la misma fila
      class_values <- unlist(strsplit(as.character(class_row), ";"))
      metabolic_domain <- class_values[2]
      metabolic_subdomain <- gsub("BRITE hierarchy", "", class_values[3])
    }

    # Crear un dataframe con los datos procesados
    subset_data <- subset(data, ONTOLOGY == relation)
    subset_data$metabolic_domain <- metabolic_domain
    subset_data$metabolic_subdomain <- metabolic_subdomain
    processed_data <- rbind(processed_data, subset_data)
  }

  return(processed_data)
}
```

**Supplementary Code 1: R script function to get hierarchical order of KEGG Pathways by requesting to genome.jp database**

```
ancestors_gene_ontologies <- function(ontologies, groups) {
  # Definir una función para obtener los ancestros de una ontología
  get_ancestors <- function(ontology) {
    url <-
sprintf("https://www.ebi.ac.uk/QuickGO/services/ontology/go/terms/%s/ance
stors?relations=is_a%%2Cpart_of%%2Coccurs_in%%2Cregulates",
                  URLencode(ontology, reserved = TRUE))
    response <- content(GET(url, accept("application/json")), "parsed")
    if (!response$results[[1]]$isObsolete) {
      ancestors <- response$results[[1]]$ancestors
      return(setdiff(ancestors, ontology))
    } else {
      return(NULL)
    }
  }

  # Obtener los ancestros para todas las ontologías de manera paralela
  plan(multisession)
  ancestors <- future_lapply(ontologies, get_ancestors)

  # Filtrar y combinar los resultados
  data <- data.frame(Group = groups, Ontology = ontologies, Ancestors =
sapply(ancestors, toString))
  data <- data[!is.null(data$Ancestors), ]

  # Escribir los resultados en un archivo CSV
  write.csv(data, "go_gene_ontologies.csv", row.names = FALSE)
}
```

**Supplementary Code 2: R script function to get ancestors of a Gene Ontology by requesting to Quick Go API.**

```
# Define function to retrieve children of a GO term
get_children_quickgo <- function(go_id) {
  # Construct URL to retrieve children of a GO term
  base_url <- "https://www.ebi.ac.uk/QuickGO/services/ontology/go/terms/"
  url <- paste0(base_url, go_id, "/children")
  response <- httr::GET(url)
  if (httr::http_type(response) == "application/json") {
    children <- httr::content(response, "parsed")
    return(children)
  } else {
    stop("Error: The response is not in JSON format.")
  }
}

# Define function to retrieve information of children of a GO term
get_children_info <- function(go_term) {
  children_quickgo <- get_children_quickgo(go_term)
  children_list <- children_quickgo$results
  children_df <- data.frame(id = character(), name = character(), stringsAsFactors = FALSE)
  for (child_info in children_list[[1]]$children) {
    child_id <- child_info$id
    child_name <- child_info$name
    child_df <- data.frame(id = child_id, name = child_name, stringsAsFactors = FALSE)
    children_df <- rbind(children_df, child_df)
  }
  return(children_df)
}

# Define function to find the first matching ancestor
find_first_matching_ancestor <- function(ancestors) {
  for (ancestor in ancestors) {
    if (ancestor %in% children_info_mf$id) {
      return(ancestor)
    } else if (ancestor %in% children_info_bp$id) {
      return(ancestor)
    } else if (ancestor %in% children_info_cc$id) {
      return(ancestor)
    }
  }
  return(NA)
}
# Define function to get the name of the first ancestor
get_first_ancestor_name <- function(ancestor_id) {
  if (ancestor_id %in% children_info_mf$id) {
    return(children_info_mf$name[children_info_mf$id == ancestor_id])
  } else if (ancestor_id %in% children_info_bp$id) {
    return(children_info_bp$name[children_info_bp$id == ancestor_id])
  } else if (ancestor_id %in% children_info_cc$id) {
    return(children_info_cc$name[children_info_cc$id == ancestor_id])
  } else {
    return(NA)
  }
}
```

**Supplementary Code 3: R script function to get on extra level of hierarchy in gene ontologies classification.** The approach consist in comparing the gene ontologies named as children of the three main gene ontologies, already explained, with all the ancestors of each of our ontologies dataset, in order to find a coincidence, meaning that this gene ontology found in common, would be connected to our GO_id observation. First of all, by *get_children_quickgo()* the children ontologies from any ontology can be taken from the database, in this case the children of the three main levels, then *get_children_info ()* will retrieve the name and id of each of the ones found; *find_first_matching ancestor* ()is used to find a common ancestor between the ones of our observations and the ones found before from the three main levels; and *get_first_ancestor_name()* ensures that the connection is correct and add the the first ancestor to our main dataset for each observation.

```
# Define function remove_duplicate_observations before analyze_regulation
remove_duplicate_observations <- function(data) {
  # Encuentra las observaciones duplicadas por "ontology", "group" y "EA_VALUE"
  duplicated_rows <- duplicated(data[, c("ONTOLOGY", "GROUP", "EA_VALUE")]) | duplicated(data[, c("ONTOLOGY", "GROUP", "EA_VALUE")],
fromLast = TRUE)

  # Cambia GOEA_up_DOWN a "NEUTRAL" en las observaciones duplicadas
  data[duplicated_rows, "UP_DOWN"] <- "NEUTRAL"

  # Elimina las filas duplicadas basadas en "ONTOLOGY", "GROUP_1", "GROUP_2", y "EA_VALUE"
  data <- data[!duplicated(data[, c("ONTOLOGY", "GROUP_1", "GROUP_2", "EA_VALUE")]), ]

  return(data)
}
# Define function analyze_regulation
analyze_regulation <- function(data) {

  # Filtra las observaciones que tienen el mismo "ontology" y "group"
  pairs <- split(data, paste(data$ONTOLOGY, data$GROUP))

  # Inicializa un vector para almacenar las observaciones a conservar
  observations_to_keep <- numeric()

  # Itera sobre cada par de observaciones
  for (pair in pairs) {
    pair_df <- as.data.frame(pair)  # Convertir el par en un dataframe

    if (nrow(pair_df) == 2) {  # Verifica que haya exactamente dos observaciones en el par
      EA_VALUE_1 <- pair_df[["EA_VALUE"]][1]
      EA_VALUE_2 <- pair_df[["EA_VALUE"]][2]

      if (EA_VALUE_1 != EA_VALUE_2) {
        observation_to_keep <- ifelse(EA_VALUE_1 > EA_VALUE_2, rownames(pair_df)[1], rownames(pair_df)[2])
        observations_to_keep <- c(observations_to_keep, observation_to_keep)
      } else {
        # Si los EA_VALUE son iguales, conservamos solo una observación
        observations_to_keep <- c(observations_to_keep, rownames(pair_df))
      }
    } else if (nrow(pair_df) == 1) {  # Si solo hay una observación en el par
      observations_to_keep <- c(observations_to_keep, rownames(pair_df)[1])
    }
  }

  # Conserva solo las observaciones seleccionadas
  data <- data[rownames(data) %in% observations_to_keep, ]
  return(remove_duplicate_observations(data))
}
```

**Supplementary Code 4: R script functions to configure the network edges color and direction.** By this code we first, eliminate the duplicated data in different cases: when both cases of variable “KEGG_up_DOWN or GOEA_up_DOWN” have an equal score for enrichment, which means that the groups are have a neutral relationship between them, that’s done by the *remove_duplicate_observations()* ; and when one of the two cases (“UP” or “DOWN”) has a higher enrichment score than the other, so the smallest one will be deleted and the bigger one will be used, as done in the *analyze_regulation()* function.
